# Structural characterization of two full length γδ TCR/CD3 complexes

**DOI:** 10.1101/2024.07.19.604350

**Authors:** Mohammed Hoque, John Benji Grigg, Trudy Ramlall, Jennifer Jones, Luke L. McGoldrick, William C. Olson, John C. Lin, Eric Smith, Matthew C. Franklin, Tong Zhang, Kei Saotome

**Affiliations:** Regeneron Pharmaceuticals, Inc. Tarrytown, NY 10591

**Author notes:** Correspondence (M.H.); (T.Z.); (K.S.).

## Abstract

The T-cell receptor (TCR)/CD3 complex plays an essential role in the immune response and is a key player in cancer immunotherapies. There are two classes of TCR/CD3 complexes, defined by their TCR chain usage (αβ or γδ). Recently reported structures have revealed the organization of the αβ TCR/CD3 complex, but similar studies regarding the γδ TCR/CD3 complex have lagged behind. Here, we report cryoelectron microscopy (cryoEM) structural analysis of two full-length γδ TCRs, G115 (Vγ9Vδ2) and 9C2 (Vγ5Vδ1), in complex with CD3 subunits. Our results show that the overall subunit organization of the γδ TCR-CD3 complexes is similar to αβ TCRs. However, both γδ TCRs display highly mobile extracellular domains (ECDs), unlike αβ TCRs, which have TCR ECDs that are rigidly coupled to its transmembrane (TM) domains. We corroborate this finding in cells by demonstrating that a γδ T-cell specific antibody can bind a site that would be inaccessible in the more rigid αβ TCR/CD3 complex. Furthermore, we observed that a Vγ5Vδ1 complex forms a TCR γ5-chain mediated dimeric species whereby two TCR/CD3 complexes are assembled. Collectively, these data shed light on γδ TCR/CD3 complex formation and may aid the design of γδ TCR-based therapies.

## Introduction

T-cells are specialized immune cells that protect the body against pathogenic threats by recognizing foreign antigens via their T-cell receptor (TCR). T-cell subsets are broadly classified based on their TCR chain usage, αβ or γδ (*1*). In humans, γδ T-cells are further subdivided by their δ-chain usage. In the peripheral blood, the most abundant γδ T-cell population utilizes the semi-invariant γ9δ2 TCR chain pairing(*2–5*). In some peripheral tissues, like the liver, intestines, and the dermis of the skin, δ1-chain expressing T-cells are enriched(*5*). Unlike αβ TCRs that almost always recognize peptides in the context of MHC/HLA, γδ TCRs can detect an array of antigens in an MHC-independent manner(*6*). While many of the ligands recognized by γδ T-cells remain unknown, the γδ TCR is thought to recognize molecules that are upregulated during cellular stress, such as lipid species in the context of CD1, and phosphonate accumulation sensed by butyrophilins(*6–9*). MHC-independent target cell recognition makes γδ T-cells attractive for cell therapies as there may be a lower risk of graft-versus-host disease(*10–12*).

On the T-cell surface, TCR chain heterodimers (TCRαβ or TCRγδ) associate with three CD3 homo- and hetero-dimers (CD3ζζ, CD3εγ, CD3εδ in humans)(*13, 14*) to form heterooctameric TCR/CD3 complexes. The TCR chains direct ligand specificity by binding directly to antigen, but unlike other receptors, lack intracellular signaling domains. On the other hand, the CD3 chains do not play a role in ligand recognition but contain ITAM sites that are phosphorylated upon ligand engagement(*15–17*). Thus, in a functional TCR/CD3 complex the TCR chains guide specific ligand engagement, while the CD3 chains drive the cellular response.

From a structural perspective, the αβ TCR heterodimer extracellular domains (ECDs) and their complexes with MHC have been studied extensively, yielding important insights into specific ligand recognition(*18–20*). More recent studies of the entire αβ TCR/CD3 complex, both with and without ligand, have provided additional insight into the organization of the TCR/CD3 complex(*21–24*). These studies demonstrated that the αβ TCR ECD, which is responsible for MHC antigen binding, is rigidly coupled to the remainder of the signaling complex via extensive contacts between TCR constant regions and CD3 subunits. Structural studies of sγδ TCR have lagged behind that of the αβ TCR and, have almost exclusively been limited to structures of TCR ECDs and their complexes with antigen(*25–27*). It has been proposed that the γδ TCR/CD3 complex may adopt an alternative organization relative to the αβ TCR/CD3, due to differences in glycosylation patterns of CD3δ and the proximity of TCR and CD3 chains as determined by crosslinking experiments(*28–31*). However, recently reported structures of γδ TCR/CD3 complexes have called these claims into question(*27*).

Here, we present cryoEM structures of two γδ TCR clones, G115 (Vγ9Vδ2) and 9C2 (γ5δ1) as full-length complexes with CD3 and in the presence of Fab fragments of antibodies targeting CD3 or TCRδ(*32, 33*). Our results show that the overall subunit organization of γδ TCR/CD3 is analogous to the αβ TCR/CD3 complex, with a conserved TM domain architecture driving assembly. However, we find that γδ TCR ECD is highly mobile due to lack of stabilizing interactions with the CD3 ECDs. This differs notably from the αβ TCR ECD, which has a relatively rigid association with the CD3 ECD that restricts its movement. Surprisingly, we observe a Vγ5 mediated dimeric species for 9C2 TCR. Collectively, our results elucidate the organization of γδ TCR/CD3 complexes and provide potential insights into how the unique structural properties γδ TCRs may underpin their specialized roles in T-cell mediated immunity.

## Results

### Purification of two full length γδ TCR/CD3 complexes

We chose two γδ TCR clones, G115 and 9C2, for which crystal structures of the ECDs have been solved previously, for structural analysis as full-length complexes with CD3(*25, 26*). The G115 γδ TCR heterodimer clone utilizes a γ9δ2 chain pairing and binds butyrophilin molecules that are upregulated upon the accumulation of intracellular phosphonate species(*34–36*). The 9C2 γδ TCR heterodimer utilizes a γ5δ1 chain pairing and directly recognizes lipid antigens presented in the context of CD1 molecules(*26*) (**Fig. S1A**). Both TCR clones were produced utilizing the same constant regions for the TCR γ-chain (TRGC1), as well as the TCR δ-chain (TRDC). We adapted a previously described scheme for producing detergent-solubilized αβ TCR/CD3 complexes(*23*) and purified both γδ TCR/CD3 complexes as singular monodisperse peaks with similar elution volumes via size exclusion chromatography (**Fig. S1B**).

### CryoEM structure of the full length G115 (Vγ9Vδ2) TCR/CD3 complex

We began by using single particle cryoEM to characterize the G115 γδ TCR/CD3 complex (henceforth referred to as G115 TCR), which uses Vγ9 and Vδ2 in its TCR chains. To aid in cryoEM image processing by increasing the molecular mass of the complex, we pre-incubated the sample with purified Fabs of the commercially available CD3ε-binding antibody OKT3(*33, 37*). The OKT3 monoclonal antibody is widely used as a T-cell activator(*38*) and was the first clinically approved monoclonal antibody(*39*). We determined a cryoEM structure of the OKT3 Fab-bound G115 TCR to 3.3 Å resolution (**Fig. 1; Fig. S2A, B; Table 1**). The cryoEM map showed density for the TM helices of the γδ TCR, the ECD and TM domains of the CD3ε/δ/γ/ζ chains, as well as two copies of the OKT3 Fab bound to the CD3ε subunits in the CD3εδ and CD3εγ dimers, respectively (**Fig. 1A and Fig. S2C**). Side chain densities were clearly resolved in most of these regions, allowing confident model building. N-linked glycans were modeled for residues N38 and N74 on the CD3δ chains, as well as N52 on the CD3γ chain (**Fig. 1B**). In the TM region, three densities corresponding to lipids positioned between the TCRγ, TCRδ, CD3ζ, and CD3γ chains (**Fig. S3A**) were apparent. We tentatively assigned cholesterol (CLR) to the interior lipid density in the lower half of the TM domain (**Fig. 1B**). In addition, we observed a density that corresponds to an S-palmityl moiety at TCRγ residue C279 **(Fig. S2C**). This residue has been predicted to be a high confidence target for palmitoylation by the SwissPalm server(*40*). However, the functional significance of this post translational modification is unknown.

**Figure 1.**
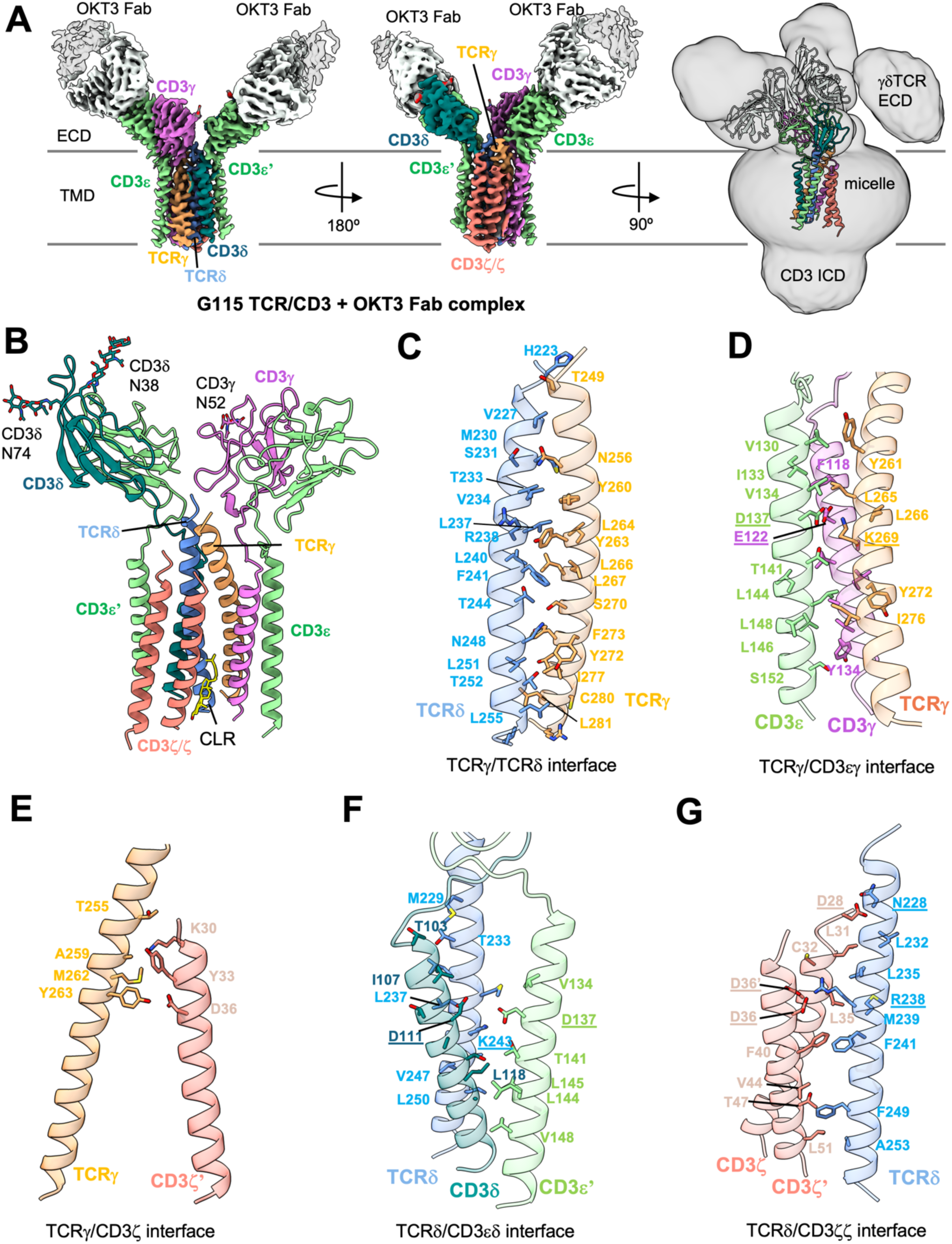
CryoEM reconstruction and modeling of the G115 (Vγ9Vδ2) TCR/CD3 complex. A) Left and middle: two views of a 3.27 Å resolution map of G115 TCR/CD3 complex bound to OKT3 fab. Right: cryoEM map was gaussian filtered to 2.5 σ and the contour level was reduced to allow visualization of weak γδ TCR ECD density. B) Structure of the G115 TCR/CD3 complex. C-G), interfaces between TCRγ/TCRδ (C), TCRγ/CD3εγ (D), TCRγ/CD3ζ (E), TCRδ/CD3εδ (F), and TCRδ/CD3ζζ (G). Chains are shown in cartoon representation and color-coded as indicated and interfacial residues are shown as sticks. Underlined residues highlight conserved electrostatic interactions between αβ and γδ TCR-CD3 interaction networks.

**Table 1.** CryoEM data, structure refinement, and validation.

|  | G115 TCR/CD3/<br>OKT3 Fab | G115 TCR<br>ECD/<br>Fab1 | 9C2 TCR ECD/<br>Fab 2 | 9C2 TCR ECD/<br>Fab 3 |
| --- | --- | --- | --- | --- |
| **Data collection and processing** |  |  |  |  |
| Magnification | 105,000 | 105,000 | 105,000 | 105,000 |
| Voltage (kV) | 300 | 300 | 300 | 300 |
| Electron exposure (e<sup>-</sup>/Å<sup>2</sup>) | ~50 | ~40 | ~40 | ~40 |
| Defocus range | -1.0 to -2.2 | -1.0 to -2.2 | -1.0 to -2.2 | -1.0 to -2.2 |
| Pixel size (Å) | 0.839 | 0.839 | 0.839 | 0.839 |
| # of Movies | 15,639 | 8,599 | 8,037 | 5,854 |
| Initial number of particles | 7.2M | 10.0M | 5.1M* | 3.1M |
| Particles selected after 2D classification | 1.7M | 250K | 64K | 46K |
| Final selected particles | 290,758 | 156,210 | 31,918 | 29,351 |
| Symmetry imposed | C1 | C1 | C2 | C2 |
| Map resolution (Å) | 3.27 | 3.21 | 3.45 | 3.46 |
| FSC threshold | 0.143 | 0.143 | 0.143 | 0.143 |
| Refinement |  |  |  |  |
| Initial Model used | 8ES7 | 8DFW | 4LFH | 4LFH |
| **Model composition** |  |  |  |  |
| Non-hydrogen atoms | 8,397 | 5,222 | 10,480 | 10,522 |
| Protein residues | 1,054 | 660 | 1,306 | 1,310 |
| Ligands | 7 | 4 | 12 | 10 |
| **R.m.s. deviations** |  |  |  |  |
| Bond lengths (Å) | 0.002 | 0.003 | 0.002 | 0.003 |
| Bond angles (°) | 0.685 | 0.605 | 0.599 | 0.596 |
| **Validation** |  |  |  |  |
| MolProbity score | 2.04 | 1.99 | 2.36 | 2.55 |
| Rotamer outliers (%) | 3.38 | 0.00 | 4.18 | 6.41 |
| Clash score | 6.55 | 12.25 | 9.75 | 11.43 |
| Ramachandran plot |  |  |  |  |
| Favored (%) | 95.92 | 94.17 | 94.57 | 94.74 |
| Allowed (%) | 4.08 | 5.83 | 5.43 | 5.26 |
| Disallowed (%) | 0.00 | 0.00 | 0.00 | 0.00 |
| **Deposition ID** |  |  |  |  |
| PDB | XXXX | XXXX | XXXX | XXXX |
| EMDB | XXXX | XXXX | XXXX | XXXX |

\* Combined particle counts from 2D template based and Topaz particle picking. Duplicate particles were removed after 2D classification.

Surprisingly, our reconstruction lacked clear density corresponding to the ECD of γδ TCR chains, suggesting it is highly mobile relative to the rest of the complex and thus not resolved upon averaging. Notably, the individual particles that contributed to our final reconstruction appear to show signal corresponding to the γδ TCR-chain ECDs, indicating the complex is intact on the cryoEM grid, and the absence of signal for the TCR ECD in the reconstruction is not due to proteolytic cleavage or degradation of the detergent-solubilized complex (**Fig. S2D**). Indeed, when we applied a gaussian filter and lowered the contour level of our high-resolution map, we see a lobe of density that likely corresponds to the flexible ECDs of the γδ TCR chains in the TCR/CD3 complex map (**Fig. 1A, right panel**). The highly mobile nature of the γδ TCR ECDs in the context of the TCR/CD3 complex contrasts starkly with αβ TCR ECDs, which are more rigidly coupled to the TM region and thereby well-resolved in cryoEM structures(*23, 24, 41*).

### Structural organization of the G115 TCR/CD3 complex

Excepting the mobile and poorly resolved ECD of the TCR chains, our structure of G115 TCR/CD3 shows that the TCRγδ chains assemble with CD3 chains in a similar manner to TCRαβ. The TCRγ and TCRδ TM helices are adjacent to each other and positioned along the center of the assembly with the three CD3 heterodimers on the perimeter (**Fig. 1B**). Sterol-shaped lipid densities, which occupy a crevice in between TCRδ, CD3δ, and CD3ζ chains, appear to play a role in stabilizing the TM domain architecture. Interestingly, sterol coordination at this site has been shown to be involved in αβ TCR activation (*42*).

The interface between the TCRγ and TCRδ TM helices is stabilized primarily by hydrophobic interactions between the helices, as well as potential hydrogen bonds between S231 (TCRδ)/N256 (TCRγ) and T252 (TCRδ)/Y272 (TCRγ) (**Fig. 1C**). TCRγ also has a significant interface with the TM helices of CD3εγ heterodimer wherein electrostatic interaction between K269 of TCRγ and two acidic residues, E122 (CD3γ) and D137 (CD3ε) near the midpoint of the membrane appears to be critical (**Fig. 1D**). TCRγ also makes several contacts with the extracellular half of one of the CD3ζ TM helices (**Fig. 1E**). Analogous to the TCRγ/CD3εγ interface, the TCRδ contacts both chains of CD3δε heterodimer, including via a bifurcated salt bridge between K243 (TCRδ) and D137 (CD3ε)/D111 (CD3δ) (**Fig. 1F**). Finally, TCRδ uses R238 to coordinate electrostatically with the D36 residues of both protomers of the CD3ζζ homodimer (**Fig. 1G**). Notably, most of the electrostatic interactions mentioned above are conserved in the αβTCR/CD3 TM domain assembly (*41*)(**Fig. S4**).

While the overall subunit organization is conserved between γδ TCR-CD3 and αβ TCR-CD3 complexes, TCR-based structural alignment of our OKT3-Fab bound model to a published αβ TCR/CD3 complex (PDB ID: 7FJD) revealed some noteworthy differences (**Fig. 2A**). This includes a ∼20° rotation in the CD3εγ heterodimer toward the region where the TCR ECD would be in a rigidly coupled complex (**Fig. 2B**). This rotation opens a gap between the CD3εγ and CD3ε’δ heterodimer ECDs, which contact each other in αβ TCR/CD3. We did not observe any substantial differences in the TM domain organizations of αβ and γδTCRs, consistent with the conserved interactions at the subunit interfaces and overall conserved positioning of sterol-shaped lipid (**Fig. 2C**).

**Figure 2.**
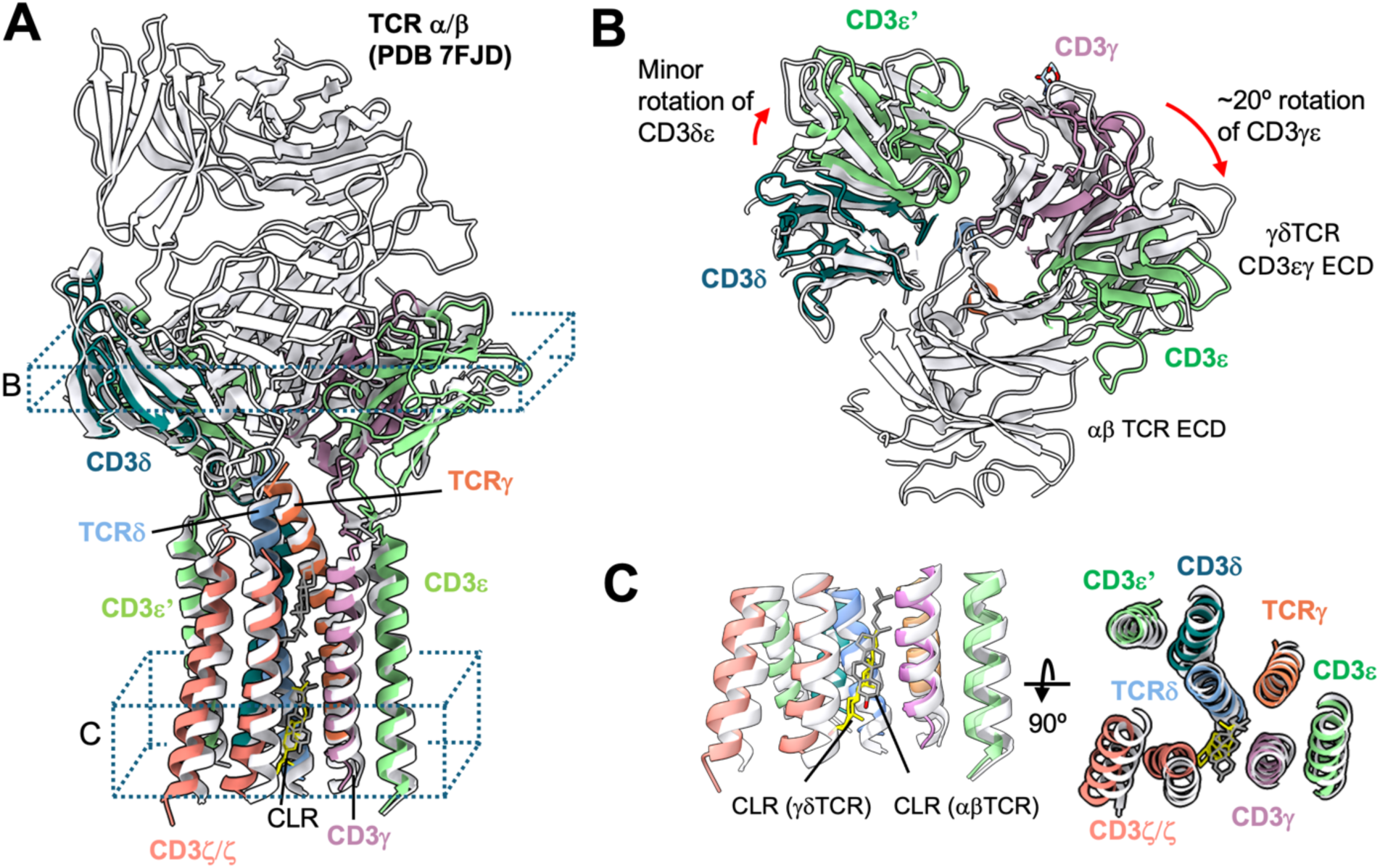
Structural comparison of the G115 TCR complex to an αβ TCR/CD3 complex. A) The G115 TCR/CD3 complex was aligned to an αβ TCR/CD3 complex (PDB:7FJD) via the TCR γ and β chains using matchmaker command in ChimeraX. G115 TCR chains are colored as indicated, while the entire αβ TCR/CD3 complex is colored white. B, C) Cross sectional and top-down views are depicted from regions indicated in A. The rotation angles shown in B were estimated between the centroids of each domain (CD3εδ or CD3εγ ECDs, comparing γδ and αβ TCRs) calculated in ChimeraX.

### CryoEM structures of mobile γδ TCR ECDs

As described above, we could only resolve weak density for the G115 TCR ECD in our OKT3 Fab complex, presumably due to TCR ECD flexibility relative to the remainder of the complex. To aid in visualization of the γδ TCR ECD by cryoEM, we made a complex of G115 TCR with the Fab fragment of a previously described TCR δ2-chain binder(*32*) (Fab 1). Unlike the OKT3-bound sample, 2D class averages for the G115 TCR/Fab 1 complex displayed well resolved features for the TCR ECD bound by Fab 1 and faint signal for the micelle/TM domain (**Fig. S5A**). We ultimately obtained a 3.2 Å resolution map in which the TCR ECD and Fab 1 V region were sufficiently resolved to permit model building (**Fig. 3A**; **Fig. S5A, B, C**; **Table 1**). We note the appearance of a weak micelle-like density in our G115 TCR/Fab 1 map upon applying a gaussian filter, further supporting the notion that the TCR ECD is flexible relative to the membrane bound portion of the molecule (**Fig. 3A, left panel**). The flexibility of the γδ TCR ECD can be rationalized by lack of sequence conservation between the γδ and αβ TCR ECD constant regions (**Fig. S4**), which apparently eliminates most of the important TCR/CD3 ECD contacts observed in αβ TCR. Our structure of G115 TCR ECD/Fab 1 complex demonstrates that the TCR ECD remains structurally intact in our detergent-solubilized cryoEM sample despite its lack of rigid association with the remainder of the complex.

**Figure 3.**
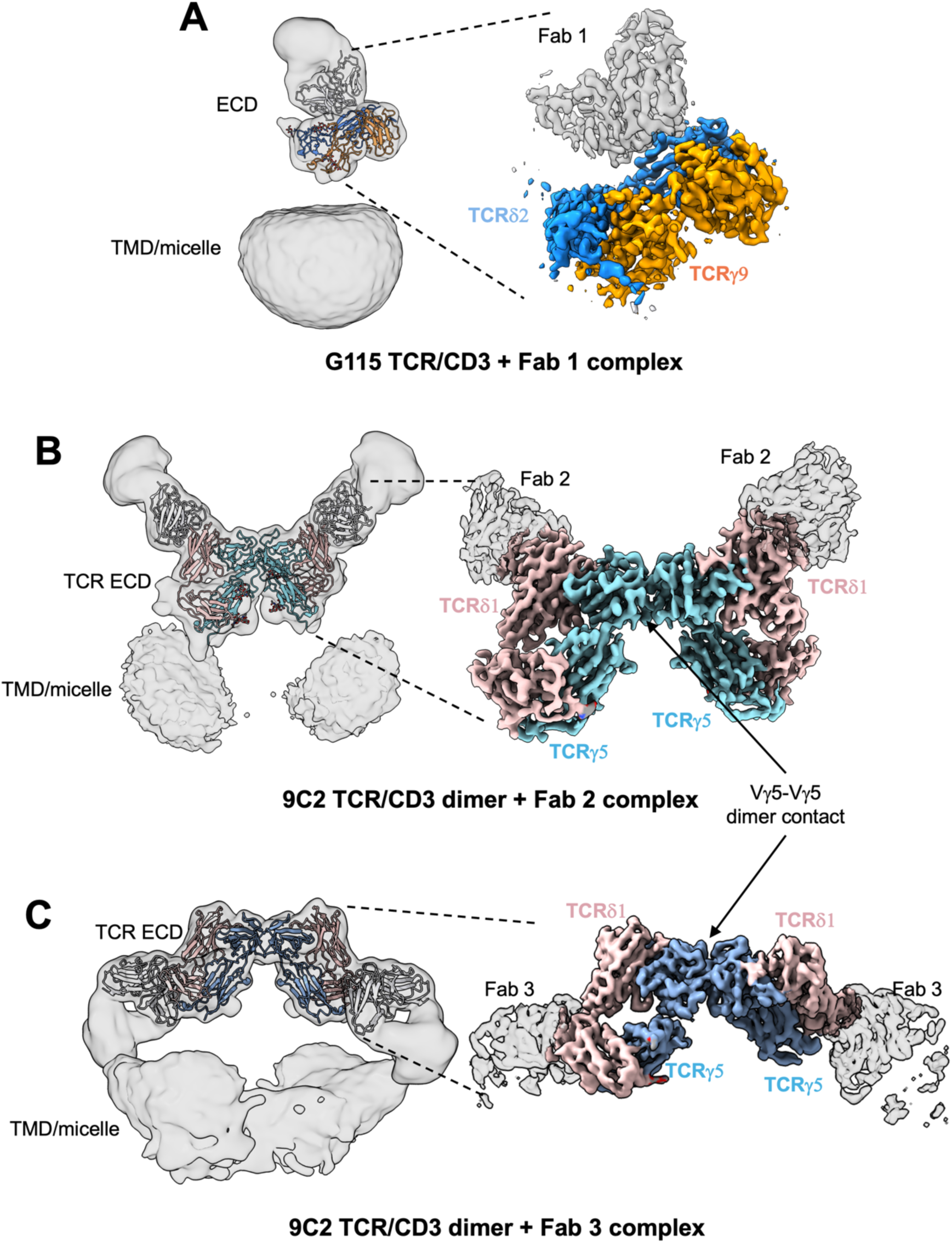
CryoEM structures of γδ TCR ECDs bound by anti-TCRVδ Fabs. A) CryoEM map of G115 TCR bound by Fab 1. B) CryoEM map of 9C2 TCR bound by Fab 2. C) CryoEM map of 9C2 TCR bound by Fab 3. Densities are color coded based on the built atomic model. Each left panel shows Gaussian filtered maps (2.5 σ) at low threshold with fitted atomic models to enable visualization of the TMD and micelle densities. Locally refined (A, B) or higher threshold sharpened (C) maps shown on the right display higher resolution features for the TCR ECD and bound Fab V domains.

We wanted to test whether the flexibility we observed for the TCR ECD was specific to the G115 TCR or a generalized phenomenon amongst γδ TCRs using different V genes. To this end, we sought to determine the structure of the detergent-solubilized 9C2 TCR/CD3 complex (henceforth referred to as 9C2 TCR) which utilized Vγ5Vδ1. To aid in TCR ECD reconstruction for the 9C2 TCR, we separately utilized the Fab fragments of two distinct TCR δ1-chain binding antibodies (Fabs 2 and 3) (*32*).

We first collected cryoEM data of 9C2 TCR bound by Fab 2. Like G115 TCR ECD, the 9C2 TCR showed high resolution features for the Fab-bound TCR ECD(**Fig. S6A**). However, in stark contrast to the monomeric G115 TCR, our 2D class averages of 9C2 TCR showed dimeric species (**Fig. S6A**). We generated a 3.5 Å resolution C2-symmetric map of this dimeric species in which side chain densities for most residues were observable for the TCR ECDs and Fab 2 variable regions (**Fig. 3B, right panel; Fig. S6; Table 1**). Applying a gaussian filter revealed two weak micellar densities situated beneath each protomer of the dimeric TCR ECD/Fab 2 complex (**Fig. 3B, left panel**). The dimer interface is in a germline encoded region of TCR Vγ5 (**Fig. 3B, right panel**). We expand on this interface in following section.

We next determined a 3.5 Å resolution cryoEM structure of the 9C2 TCR bound to a Fab 3 (**Fig. 3C, right panel; Fig. S7; Table 1**). In our structure, Fab 3 binds TCR δ1 at the linker connecting the variable and constant regions, employing a binding angle distinct from Fab 2. Like the 9C2/Fab 2 complex, we observed a γ5-chain mediated dimer species in the Fab 3 complex structure, with two micellar densities positioned beneath the TCR chain ECDs (**Fig. 3C, left panel)**. In addition, we observed a small population of particles (∼10K) where the dimeric γδ TCR is embedded in one micellar density (**Fig. S7D**). Unfortunately, we were unable to reconstruct a high resolution model of this set of particles, which may have been more representative of a γδ TCR dimer constrained in *cis* at the cell membrane than our current dimer structures, where the TM domains of each TCR/CD3 complex are contained within separate micelles. Taken together, these data indicate that γδ TCR ECDs are flexible across different clonotypes and reveal the presence of a γ5-mediated dimeric species of detergent-solubilized 9C2 TCR/CD3 complex.

### Structural analysis of the 9C2 TCR ECD dimer interface

Using the higher resolution 9C2 TCR/Fab 2 structure, we analyzed the residues that are involved in the γ5-mediated dimeric interface (**Fig. 4A**). R19 of each γ5 protomer makes a hydrogen bond with the backbone nitrogen of S56 at the periphery of the interface. The core of this interface is mediated by a π-π interaction between Y72 in each protomer. Additionally, hydrogen bonds were observed between side chains of residues D60/R86 and H74/S21, while S58 makes a hydrogen bond with the backbone nitrogen of G69. To assess the sequence conservation of Vγ5 residues at the dimer interface, we performed a sequence alignment of TCR γ-chain variable regions (TRGV) (**Fig. 4B**). Overall, the variable region is well conserved throughout all the sequences we analyzed apart from TRGV9, which is exemplified by the G115 TCR that only showed monomeric species in this study. We find that the R19/S56 interaction is likely only found in the TRGV5-utilizing TCRs, as the arginine in position 19 is a glycine in TRGV2/3/4/8. Likewise, the D60/R86 electrostatic interaction is not conserved in other TCRγ chains; TRGV3, the only other TRGV in which D60 is conserved, has a glutamine at position 86. Y72, which makes an aromatic interaction at the core of this dimeric interface, is conserved in TRGV2 and TRGV3 but replaced with charged residues in TRGV4, V8, and V9. Overall, the residues mediating Vγ5 dimerization are not well conserved, but we cannot rule out the possibility of different modes of dimerization used by other TCRγ chains.

**Figure 4.**
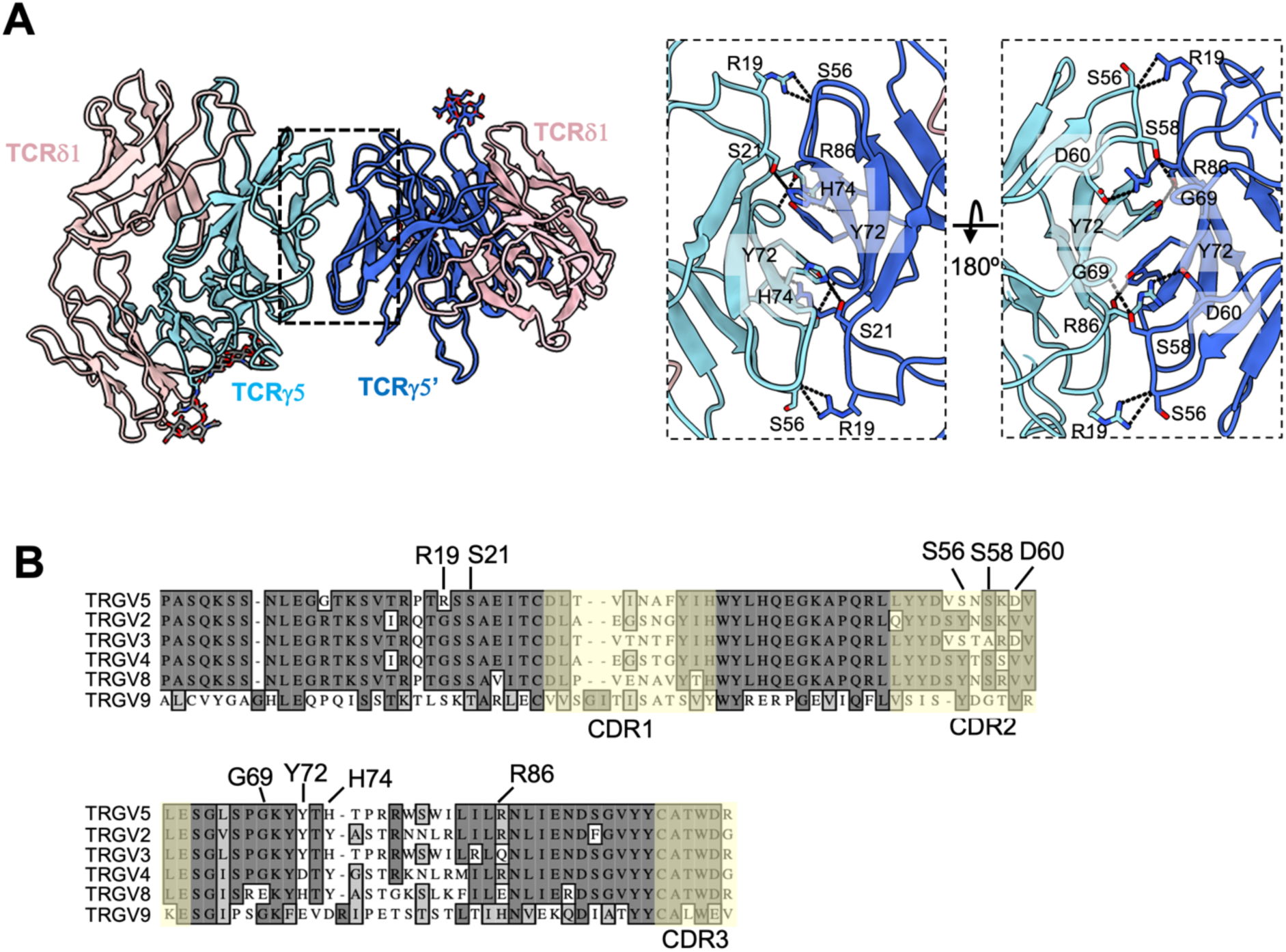
Vγ5Vδ1 TCR dimer interface and its sequence conservation. A) Structure of 9C2 TCR ECD (Fab 2 complex used, Fab models hidden for clarity) with each TCRγ protomer colored a different shade of blue. Analysis of the TCRVγ5-mediated dimer interface with interacting residues shown as sticks and dashed lines representing putative hydrogen bonds. Note: TCRγ Y72 of protomer 1 forms a π-π interaction with Y72 of protomer 2. B) Sequence alignment of various germline encoded TCR γ-chain variable regions. Residues that have been identified in the dimerization interface are labeled. CDRs are indicated with a faint yellow tint.

### *Fab 3 binds a TCR*δ*1* epitope that would be obscured in the αβ TCR-CD3 complex

We noted that Fabs 2 and 3 bind the 9C2 TCRδ1 chain using distinctly different mechanisms (**Fig. 3B and C**). While Fab 2 approaches the TCRδ V region from the opposite side of the weak micellar densities (**Fig. 3B, left panel**), Fab 3 binds using an angle such that its C region is positioned adjacent to the micelles (**Fig. 3C, left panel**). Considering this unique binding mode that apparently brings Fab 3 in potential overlap with the membrane plane, we surmised that the binding of Fab 3 to the 9C2 γδ TCR/CD3 complex is only possible due to the lack of rigid coupling between the γδ TCR ECD and the rest of the complex. Indeed, structural alignment of our γδ TCR ECD/Fab 3 complex to αβ TCR/CD3 complex indicated that the region where Fab 3 would bind onto TCRα is occluded by the CD3δ ECD of the CD3εδ heterodimer (**Fig. 5B**). To confirm that Fab 3 can bind a Vδ1 γδ TCR/CD3 complex expressed on cell surface, we tested the ability of the parental IgG of Fab 3 (Ab 3) to expand Vδ1 γδ T cells from human donor PBMCs. Plate-bound Ab 3 indeed selectively expanded Vδ1 γδ T-cells after 13 days of culture (**Fig. 5C,D**). This data supports our structural observations by showing that the γδ TCR ECD is highly flexible in a functional cellular environment, as the Fab 3 epitope would be masked in the more rigid αβ TCR/CD3 complex.

**Figure 5.**
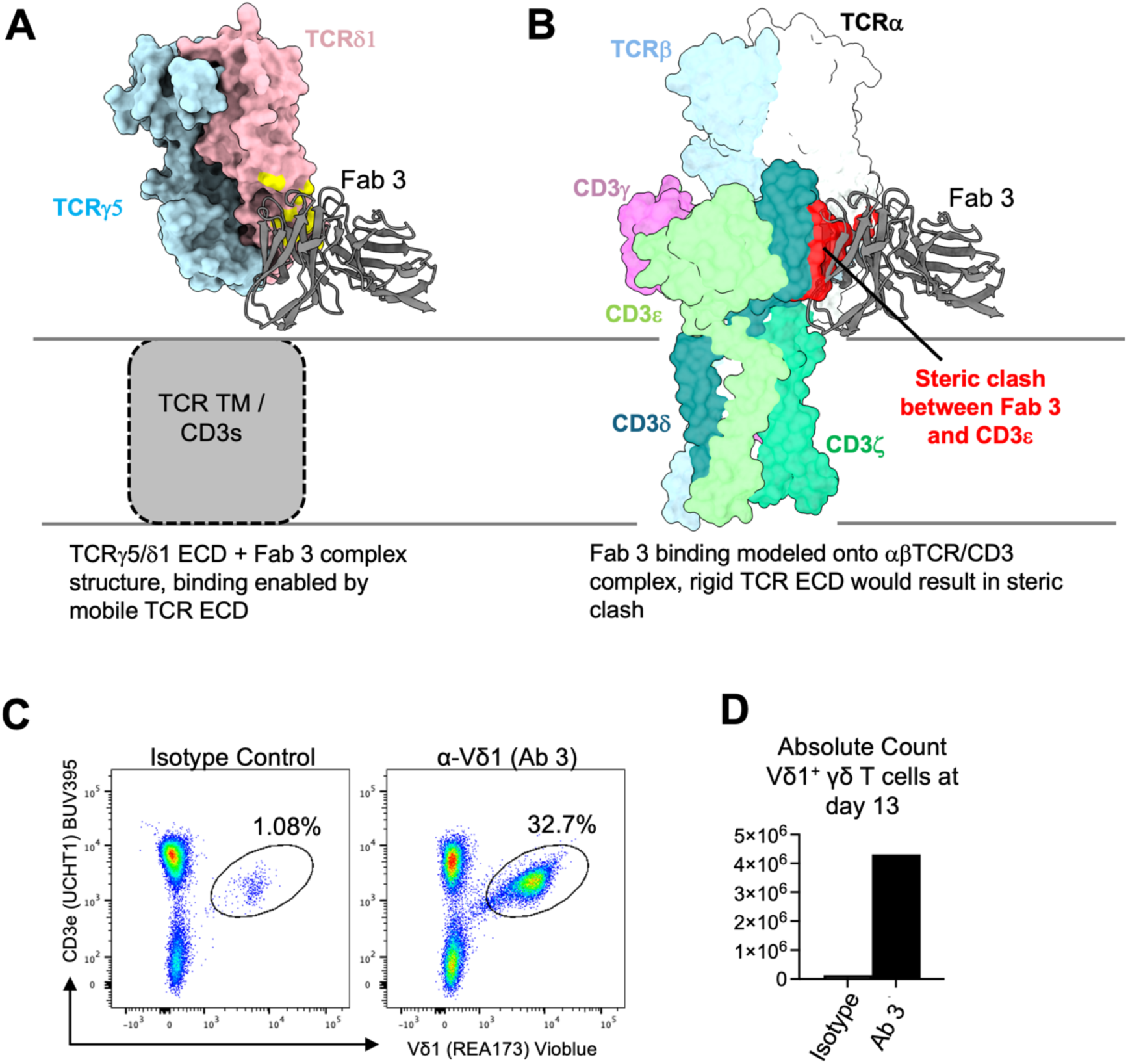
Fab 3 binding region is masked in the rigid αβ TCR/CD3 complex. A) Depiction of the Fab 3 binding site in a Fab 3 bound 9C2 γδTCR ECD structure. The Fab variable region is shown as cartoons, whereas the TCR ECD is shown as surface. The Fab 3 epitope is highlighted in yellow. B) Bound Fab 3 was modeled onto the αβ TCR from PDB ID 8ES7 through alignment of the TCR β-chain to TCRγ. The Fab variable region is shown as a cartoon, whereas the TCR ECD is shown as surface view. Apparent steric clash between Fab 3 and CD3ε is highlighted in red. C, D) Expansion of Vδ1 γδ T-cells from human donor PBMCs following culture with Ab 3. T-cell purity (C, gated on live, CD45+ cells), and counts (D) were assessed on day 13.

## Discussion

We utilized cryoEM to determine the structures of two clonotypic γδ TCR/CD3 complexes bound by the Fab fragments of antibodies directed against CD3 or TCR δ-chain. We find some overall structural similarities between the αβ and γδ TCR/CD3 complexes (**Fig. 2**). Both TCRs utilize the same stoichiometry in their CD3 chain usage **(Fig. 1**), retain a similar transmembrane domain architecture (**Fig. 2)**, and bind a lipid resembling cholesterol at analogous locations **(Fig. 2C**)(*23, 24, 41*). In addition, our results show that the γδ TCR/CD3 complex differs in two major ways from αβ TCR/CD3. First, the extracellular domain of the γδ TCR heterodimer is flexible relative to the membrane embedded portion of the molecule and the CD3 ECD heterodimers (**Figs. 1, 3, 5)**, due to absence of the extensive interactions between the γδ TCR constant regions and the CD3 ECDs that are present in αβTCRs. Secondly, for the 9C2 (Vγ5Vδ1) TCR, we observed a surprising dimeric species mediated by the γ5 chain (**Fig. 3B, C**).

The flexibly coupled ECD of γδ TCR/CD3 may allow for binding to a more diverse range of antigens than αβ TCRs, which are restricted to pMHCs and related molecules (**Fig. S8**). In the case of the αβ TCR/pMHC-I interaction, the TCR typically engages its ligand in a head-to-head orientation using an overall conserved docking polarity that places the TCR Vα and Vβ over the α2 and α1 helix of MHC-I, respectively(*43*). This docking polarity has been demonstrated to be critical for productive TCR signaling and coreceptor localization with CD3ζ(44). As such, a rigidly coupled αβ TCR ECD would be required to effectively transmit the docking polarity at the TCR/pMHC interface to the transmembrane and cytosolic side of the TCR/CD3/coreceptor complex, where signaling occurs. However, for γδ TCR, CD4/CD8 co-receptors are not required (*45*), and there is not only a wider range of ligands that can be engaged but the mode of engagement can vary. For example, the 9C2 TCR binds CD1d/lipid complexes in a classical MHC-like interaction(*26*), while the G115 TCR recognizes the B7-molecule BTN2A1 in a side-on orientation that leaves the CDRs largely uninvolved(*36*). This difference in ligand engagement geometry is further highlighted when considering the metabolite presenting MHC-like molecule MR1. Although αβ TCR engages MR1 in MHC-like fashion by docking on top of the metabolite-binding cleft(*46*), γδ TCRs displays diversity in its engagement of MR1; G83 (Vγ8Vδ3) TCR binds the side of the metabolite binding cleft in MR1 independent of antigen(*47*) and the G7 (Vγ9Vδ1) TCR clone binds distal from the metabolite binding cleft(*48*). The latter non-canonical γδ TCR/MR1 interaction presumably requires a flexible γδ TCR ECD, as observed in our cryoEM data, to contact the distal portion of the MR1 molecule.

In addition to diverse ligand recognition, the flexibility of γδ TCRs may explain the differential glycosylation patterns observed on CD3δ in αβ and γδ TCR/CD3 complexes (*28*). A highly flexible γδTCR ECD may allow increased access for glycosyltransferase enzymes to the surface of CD3δ, resulting in the addition of more complex N-linked glycans in the case of γδTCR/CD3. Furthermore, a flexible TCR ECD provides insight into why the γδ TCR lacks mechanosensitivity relative to αβ TCRs (*49*). The absence of rigid coupling between γδ TCR ECD and the CD3 subunits and TM region likely prevents the mechanical force produced by antigen binding to be transduced efficiently to the membrane and signaling components.

We report a surprising finding that the 9C2 TCR/CD3 complex displays Vγ5-mediated dimerization (**Figs. 3 and 4**), while our cryoEM data for G115 (Vγ9Vδ1) TCR/CD3 complex only showed monomeric species (**Fig. 3A**). Of note, this mode of dimerization is also present as crystal contacts in x-ray structures of 9C2, both alone and in complex with CD1d antigen (**Fig. S9**) (*26*). This suggests that dimerization may be promoted by high local concentration of the TCR ECD and does not require the full-length complex. The relevance of this dimer for γδ TCR function needs further study, but we speculate that this interface may act to induce local clustering of TCRs on the cell surface(*50, 51*) that could result in a more potent T-cell response.

During the preparation of our manuscript another group, Xin et al., independently reported structures of both the G115 and 9C2 γδ TCR chains in the context of a TCR/CD3 complex(*27*). Overall, our two studies reach similar conclusions; the ECDs of the γδ TCR chains are flexibly tethered to the TM portion of the molecule and the 9C2 TCR/CD3 complex undergoes Vγ5 chain-mediated dimerization. Notably, Xin et al. conduct elegant experiments supporting the potential cellular role of γ5 chain-mediated dimerization. We take this opportunity to note small differences between our two studies with regards to dimerization. Xin et al. report that the dimeric 9C2 TCR/CD3 complex has a left-shifted SEC elution volume relative to that of the G115 TCR/CD3 complex, while we observed that the elution volumes of 9C2 and G115 are essentially unchanged, indicating the 9C2 TCR dimers were generated upon sample concentration or vitrification. Further, a majority of our particle images of dimeric TCR ECD apparently contained two individual micelles that were attached via the γ5 chain, whereas Xin et al. report 9C2 TCR complexes dimerized within a single micelle (*27*). 2D class averaging suggested that only a minor population of particles in our samples were of this latter species. The basis for the subtle differences in dimerization behavior observed in our experiments and those of Xin et al. remain unclear. Nonetheless, our studies reinforce each other to elucidate the unique structural features of γδ TCR/CD3 complexes.

## Methods

### Construct design

TCR-CD3 construct designs were adapted from previous approaches (**Fig. S1**)(*23*). Briefly, TCR and CD3 chain DNA constructs were codon-optimized and synthesized by GenScript. The full-length G115 and 9C2 TCR constructs were comprised of the δ-chain followed by the γ-chain with an intervening furin cleavage sequence and a P2A cleavage site. The CD3 construct was designed as previously described(*23*).

### γδ TCR-CD3 expression

Protein used for TCR-CD3 complexes were expressed using BacMam-mediated viral transduction in HEK293F cells. BacMam virus for each construct was produced in ExpiSf9 cells maintained in ExpiSf CD media. P1 viral stocks were concentrated by centrifugation at 72,500 × g in a Ti45 rotor for one hour at 4°C. The viral stocks were then resuspended in 2% FBS/Freestyle 293 media. HEK293F cells were then infected with a 1:1 mixture of G115 TCR/CD3 viruses or 9C2 TCR/CD3 viruses (as shown in **Fig. S1A**) and incubated at 37°C. 12-16 hours post infection, 10 mM Sodium Butyrate was added to the culture and collected 48 hours after infection. Cells were then harvested by centrifugation, washed with ice cold PBS + Protease inhibitor, and stored at −80°C for downstream applications.

### γδ TCR/CD3 isolation and purification

HEK293F cells were thawed and resuspended in buffer containing 20 mM HEPES pH 8.0, 150 mM NaCl, 1% Glyco-diosgenin (GDN), and EDTA-free cOmplete protease inhibitors (Roche). The mixture was stirred for 1 hour at 4°C and then clarified by centrifugation at 30,000xg for 20 minutes. The lysate was then added to GFP nanobody-coupled Sepharose resin pre-equilibrated with SEC buffer (20 mM HEPES pH 8.0, 150 mM NaCl, 0.01% GDN) and rotated at 4°C for 1.5 hours. The resin was then collected in a gravity column and washed with SEC buffer several times. The washed resin was then collected and PreScission protease and 0.5 mM DTT were added and incubated overnight at 4°C to liberate the TCR-CD3 complex from the Sepharose beads. This mixture was then added to a gravity column, the flow through was collected, concentrated using a 100 MWCO Amicon Ultra centrifugal filter, and injected into a Superose 6 Increase 10/300 GL column. Peak fractions were collected, concentrated using a 100 MWCO filter, and either frozen at −80°C or prepared directly for cryoEM analyses.

### Fab and γδTCR/CD3 complex sample preparation

OKT3 Fab was cleaved from OKT3 IgG antibody using IdeS enzyme and standard protocols. Anti-TCRVδ chain Fabs 1, 2, and 3 were cleaved from IgGs using Pierce Mouse IgG1 Fab and F(ab’)2 Preparation Kit (ThermoFisher) using protocols provided by the manufacturer. Fc domains were removed from the samples using CaptureSelect multispecies Fc resin (ThermoFisher). Fabs were further purified by SEC, concentrated and stored at −80°C prior to use.

G115 TCR/CD3 + OKT3 Fab complex was generated by mixing the two components at a 1:1.2 molar ratio. G115 TCR/CD3 + Fab 1, 9C2 TCR/CD3 + Fab 2, and 9C2 TCR/CD3 + Fab 3 complexes were generated by mixing the full-length receptor and Fab at 1:0.6 molar ratio. An internal CD3-binding Fab was added to each of the latter three samples, but we were unsuccessful in obtaining high resolution reconstructions of the complex between CD3 and this Fab.

### CryoEM grid preparation and data collection

UltrAuFoil 1.2/1.3 grids, freshly plasma cleaned in a Solarus II (Gatan) using a H2/O2 gas mixture, were used for each sample. In the case of the G115 TCR/CD3 + OKT3 Fab complex, 0.01% fluorinated octyl-maltoside (FOM) was added immediately before freezing to help overcome preferred orientation (*52*). Samples were plunge frozen (blot time 7-15 seconds, blot force 0) in liquid ethane cooled by liquid nitrogen in a Vitrobot Mark IV operated at 4°C and 100% humidity.

Grids were loaded into a Titan Krios G3i electron microscope equipped with a BioQuantum K3. Images were collected in counting mode at a magnification of 105kx, yielding a pixel size of 0.839 Å. For data collection, a defocus range of −1.0 to −2.2 µm was used, the energy filter was inserted with a width of 20 eV, and the 100 µm objective aperture was inserted. Each movie was dose fractionated into 46 frames over a 1.6 second exposure and had a total dose of ∼40 or ∼50 e-/Å^2^.

### CryoEM data processing

All datasets were preprocessed in a similar manner. Briefly, movies were imported into cryoSPARC v4.3 (*53*) and preprocessed using Patch Motion Correction and Patch CTF Estimation. Micrographs were curated using a 3.5 Å CTF cutoff, except the Fab 3 bound 9C2 complex which utilized a 4.0 Å cutoff. 2D class averages generated from blob-picking random subsets of micrographs were used as templates to pick particles from the respective datasets. TOPAZ picking(*54*) was also used for the Fab 2-bound 9C2 TCR/CD3 complex. Further processing steps are described below:

#### OKT3 bound G115 TCR/CD3 complex

7.2 M particles from template-based picking were subjected to two rounds of 2D classification were conducted, only keeping particles contributing to class averages with clear features of the micelle-embedded receptor complex, yielding 1.7 million particles. Three successive cycles of ab initio and heterogenous refinement were done, with the best class selected to proceed to the next round. This resulted in a clean class of 290K particles. Non-uniform refinement(*55*) was then performed yielding a map with a nominal resolution of 3.27 Å.

#### Fab 1 bound G115 TCR/CD3 complex

10.0 M particles from template-based picking were subjected to multiple rounds of 2D classification, keeping class averages clearly showing the TCR ECD/Fab 1 complex, yielding 250K particles. Two successive cycles of ab initio and heterogenous refinement were done, with the best class selected to proceed to the next round, resulting in a final subset of 156K particles. Non-uniform refinement was then performed yielding a map with a nominal resolution of 3.58 Å. Local refinement using a mask focusing on the TCR ECD and the Fab V domain yielded a map with a nominal resolution of 3.21 Å.

#### Fab 2 bound 9C2 TCR/CD3 complex

Multiple rounds of 2D classification were conducted separately on 2D template-picked (2.5 M) and TOPAZ-picked (2.8 M, using a model generated from clean template picks from this dataset) particles, selecting class averages showing clear features of the dimerized TCR ECD/Fab 2 complex. Particles after 2D classification from 2D template picking and TOPAZ picking were combined and duplicates removed, resulting in a stack of 64K particles that were subjected to ab initio reconstruction with 3 classes. 32K particles from the best ab initio class were refined with C2 symmetry using non-uniform refinement, yielding a map with a resolution of 3.41Å. Local refinement using a mask focusing on the TCR ECD and the Fab V domain yielded a C2-symmetric 3.45 Å resolution map with improved features.

#### Fab 3 bound 9C2 TCR/CD3 complex

3.1 M particles from template picking were subjected to multiple rounds of 2D classification, keeping class averages showing dimerized TCR/Fab 3 complex with two micelle densities present. ∼46K particles from this process were then subjected to ab initio refinement (3 classes). The best class (29K particles) was selected and nonuniform refinement was performed with C2 symmetry yielding a map with a nominal resolution of 3.46 Å. We also observed a rare population of particles that displayed 2D class averages showing dimerized 9C2 ECD contained within one micellar density; however, we were unable to resolve a high-resolution model of these particles.

### Model building and refinement

The model of the G115 TCR/CD3 complex was built using a published structure of αβ TCR/CD3 complex (PDB ID: 8SE7) as an initial model. The TM domains of the αβ TCR chains were deleted and replaced with an AF2(*56*) prediction of the TCR γ- and δ-chains. The model of the G115 TCR ECD was built using a published crystal structure of the G115 TCR ECD bound to BTN2A1 (PDB ID: 8DFW) as an initial model. The 9C2 TCR model building used a published ECD crystal structure (PDB: 4LFH) as an initial model. All Fabs were modeled using AF2 predictions. The models were iteratively built manually in COOT 0.9.8.94 EL(*57*) and real-space refined in PHENIX 1.21.1(*58*). All refinements were performed with secondary structure, Ramachandran, and geometry restraints turned on. The structures were validated with Molprobity as implemented in PHENIX. All structural figures were generated in ChimeraX(*59*) or Pymol 2.5.4(*60*).

### Expansion of Vδ1 expressing γδ T-cells

Human Vδ1^+^ γδ T cells were expanded by incubating healthy donor PBMCs with plate-bound agonistic α-Vδ1(Ab 3) or Isotype antibody control in media containing recombinant human IL-2. At day 7, cell cultures were transferred to new plates without agonistic α-Vδ antibody and continued to culture in media containing IL-2. Final T cell counts, and purity were assessed on day 13 by AOPI labeling (Nexcelom) and flow cytometry using detection antibodies for human CD45 (clone HI30), human CD3 (clone UCHT1), and human Vδ1 (clone REA173).

### Multisequence alignment and annotation of TCR chains

TRDC, TRAC, TRGC1/2, TRBC1/2, and TRGV protein sequences were obtained from UniProt and aligned via MUSCLE(*61*).

## Acknowledgements

We thank Yi Zhou, Erich Joachim Goebel for their invaluable input, Nam Nguyen for assistance with cell culture and protein expression, and the Regeneron cloud/HPC teams for supporting cryoEM data storage and processing.

## Author contributions

M.H., K.S., T.Z., L.L.M, M.C.F., E.S., W.C.O, and J.C.L. conceptualized the studies. M.H., K.S., L.L.M, T.R., J.J., expressed and purified proteins. M.H. and K.S. prepared samples for cryoEM, acquired, and built the atomic models with contributions from M.C.F. J.B.G. conducted the γδ T-cell expansion and flow cytometry studies. M.C.F., E.S., W.C.O, and J.C.L. supervised the overall project. M.H., K.S., and T.Z. drafted the manuscript with contributions from M.C.F., E.S., W.C.O, and J.C.L. The manuscript was finalized by all authors.

## Data and materials availability

All data needed to evaluate the conclusions in the paper are present in the paper and/or the Supplementary Materials or have been deposited in the following databases: The cryoEM maps and coordinates of the G115-OKT3, G115-Fab1, 9C2-Fab2, and 9C2-Fab3 are deposited and available at PDB and EMDB with access codes XXX, EMD-XXX, XXX, EMD-XXX, XXX, EMD-XXX, XXX and EMD-XXX, respectively. Regeneron materials described here may be made available to qualified, academic, noncommercial researchers through a material transfer agreement upon request at https://regeneron.envisionpharma.com/vt_regeneron/. For questions about how Regeneron shares materials, use the email address.

---

**Figure S1.**
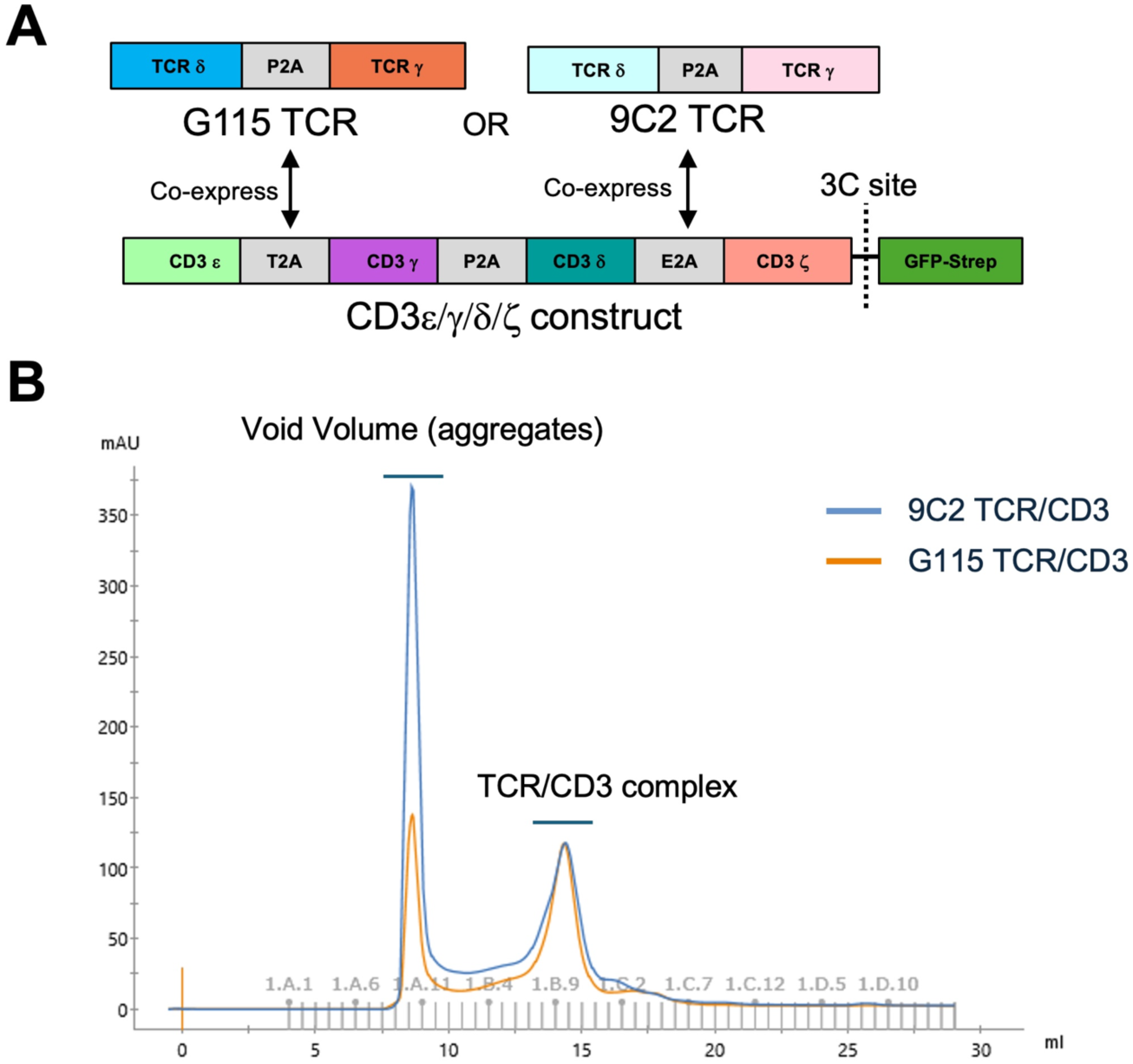
Purification of two clonotypic γδ TCR/CD3 complexes. A) Schematic representation of constructs used to express the γδ TCR/CD3 complexes. B) Size exclusion chromatography profile using Superose 6 Increase 10/300 GL column of detergent-solubilized γδ TCR/CD3 complexes following GFP-strep tag cleavage.

**Figure S2.**
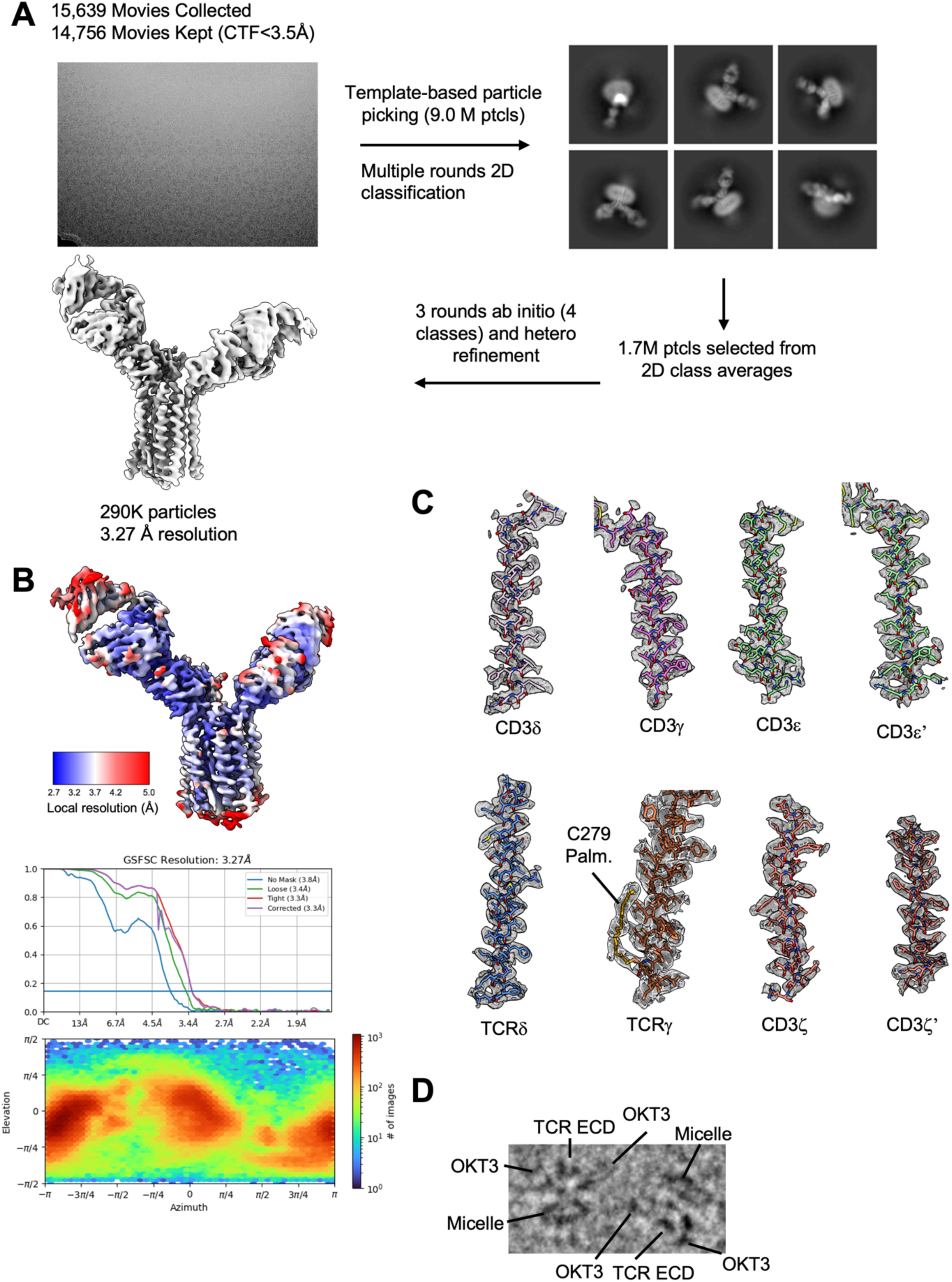
CryoEM structure of OKT3 Fab-bound G115 TCR/CD3. A) Data processing pipeline. Only representative 2D class averages are shown. B) CryoEM map filtered and colored according to local resolution, gold standard FSC curve, and particle orientation distribution as produced by CryoSPARC. C) Density fit of indicated TCR/CD3 chain TM domains into cryoEM densities. The cryoEM density protruding out of TCRγ C279 resembles a palmitoylated cysteine. This is shown for illustration but is not included in the deposited model. D) Two individual particles displaying the ECDs of the γδ TCR chains.

**Figure S3.**
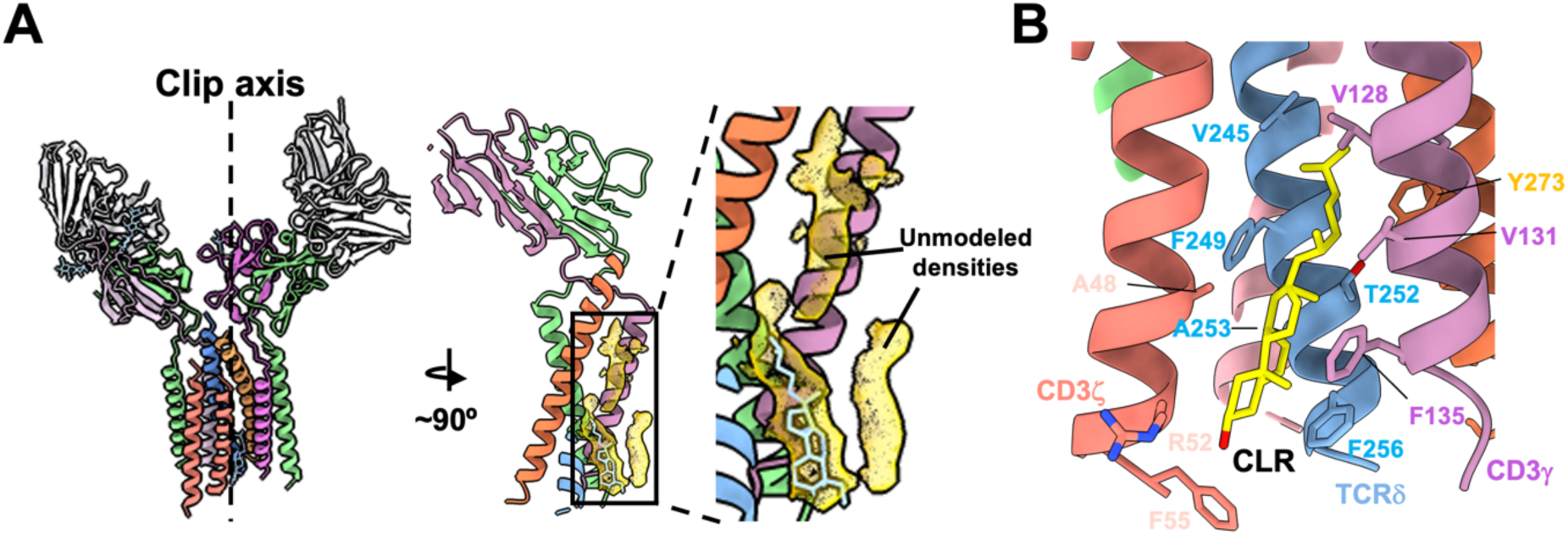
Sterol-shaped densities in γδTCR TM domain. A) Analysis of sterol like densities in the TM region of the TCR. Boxed region in the middle is enlarged in the right panel. B) van der Waals interactions between a putative cholesterol (CLR) and residues in the TCR/CD3 chains.

**Figure S4.**
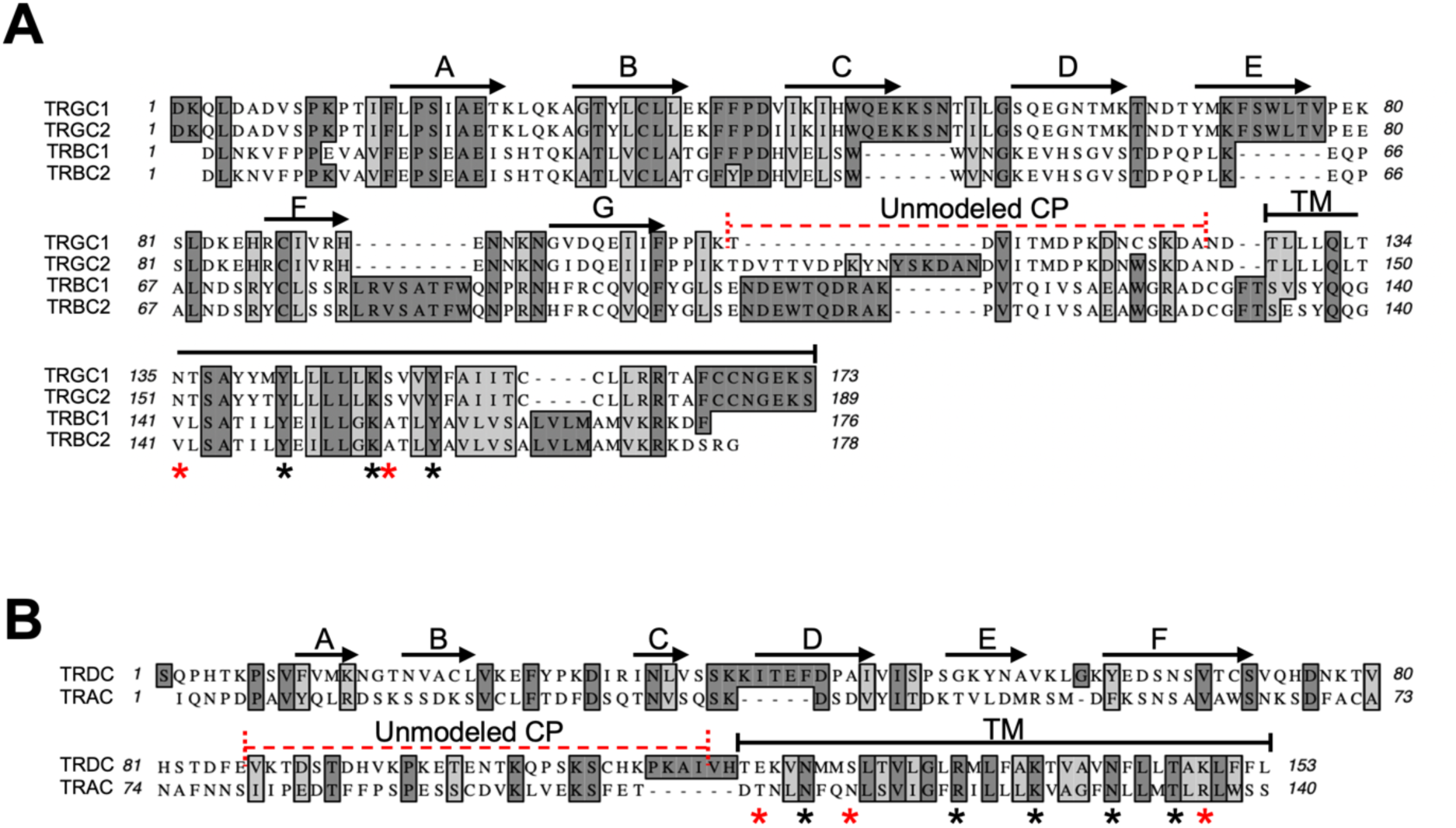
Annotated multisequence alignment of TCR constant regions. A) Multisequence alignment of TRGC1/2 and TRBC1/2 protein sequences. B) Multisequence alignment of TRDC and TRAC protein sequences. Red dashed lines represent the flexible portions of the connecting peptides that are not resolved in our ECD or TM domain-containing structures. Asterisks are used to label γδTCR residues involved in apparent hydrogen bonds with CD3. Black asterisks represent residues that are conserved between αβ and γδ TCR chains. Red asterisks represent residues that are specific to γδ TCR/CD3 complexes. Note: sequences are numbered as they appear in Uniprot, using the following accession codes: TRGC1: P0CF51; TRGC2: P03986; TRBC1: P01850; TRBC2: A0A5B9; TRDC: B7Z8K6; TRAC: P01848.

**Figure S5.**
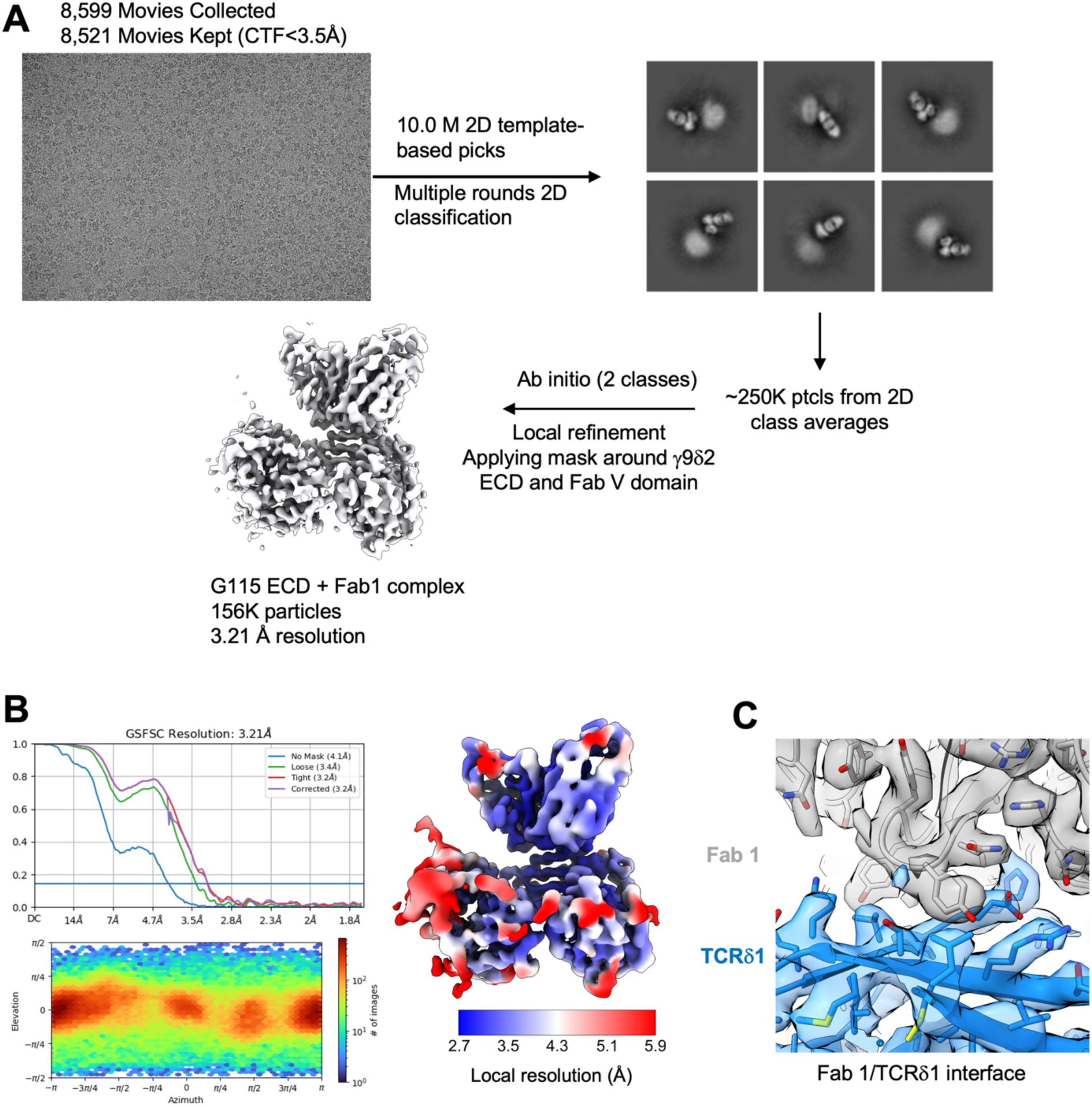
CryoEM structure of Fab 1-bound G115 TCR ECD. A) Data processing pipeline. Only representative 2D class averages are shown. B) CryoEM map filtered and colored according to local resolution, gold standard FSC curve, and particle orientation distribution as produced by CryoSPARC. C) Expanded view showing fit of model to density at the Fab 1-Vδ2 interface.

**Figure S6.**
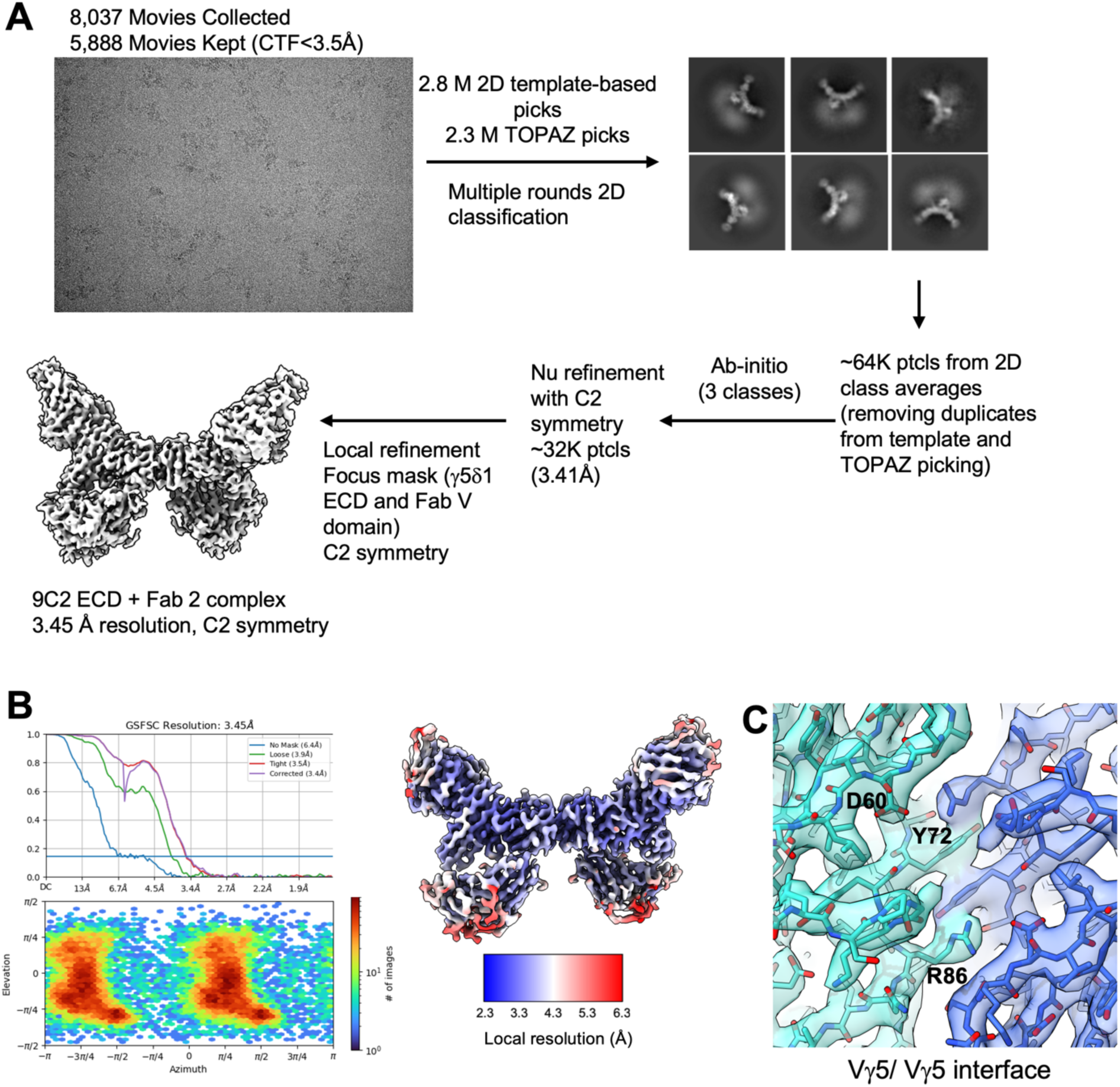
CryoEM structure of Fab 2-bound 9C2 TCR ECD. A) Data processing pipeline. Only representative 2D class averages are shown. B) CryoEM map filtered and colored according to local resolution, gold standard FSC curve, and particle orientation distribution as produced by CryoSPARC. C) Expanded view showing fit of model to density at the Vγ5/Vγ5 interface.

**Figure S7.**
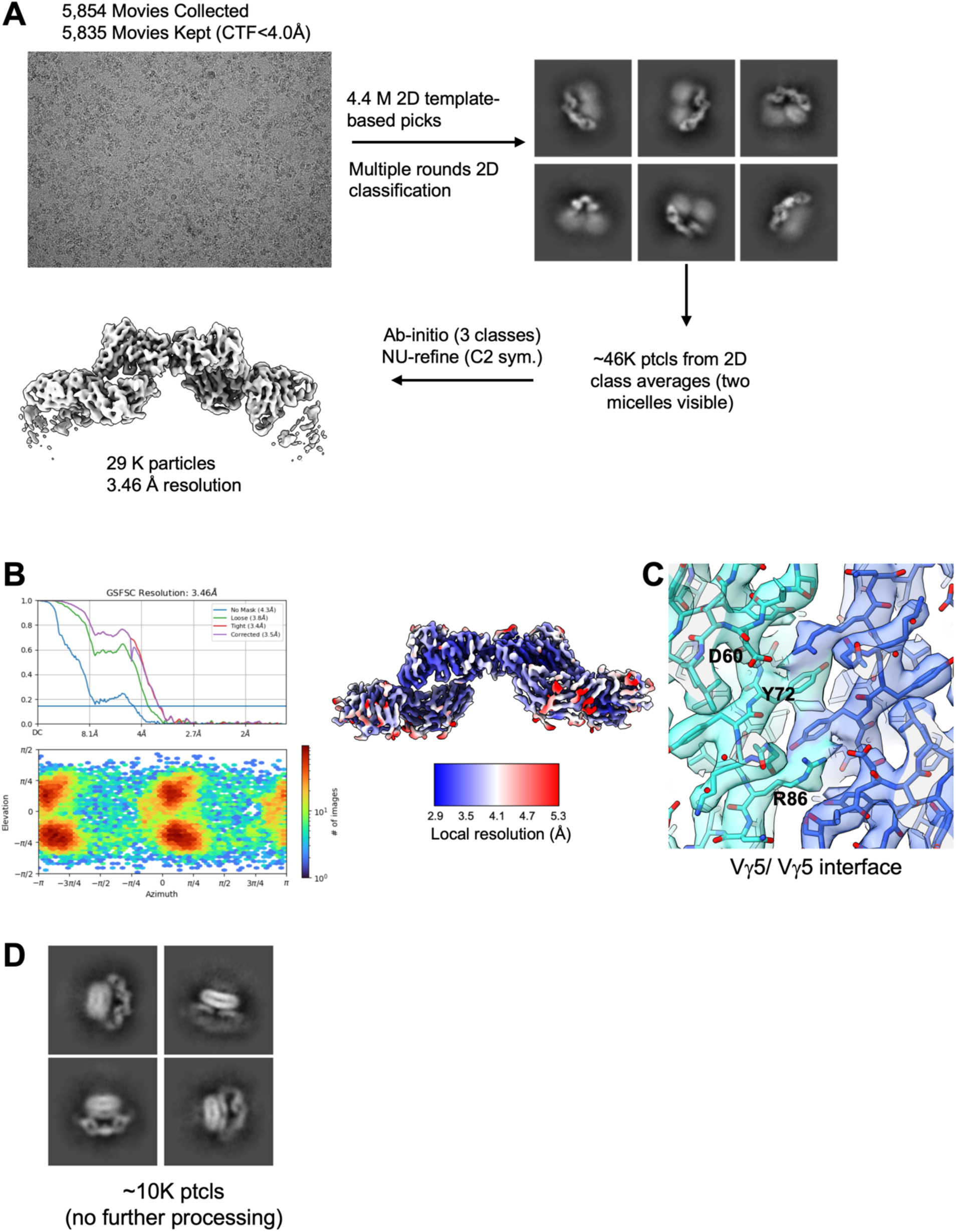
CryoEM structure of Fab 3-bound 9C2 TCR ECD. A) Data processing pipeline. Only representative 2D class averages are shown. B) CryoEM map filtered and colored according to local resolution, gold standard FSC curve, and particle orientation distribution as produced by CryoSPARC. C) Expanded view showing fit of model to density at the Vγ5/Vγ5 interface. D) 2D class averages of a minor subset of particles showing dimeric 9C2 TCR ECDs contained within one micellar density. 3D reconstruction was not successful for these particles.

**Figure S8.**
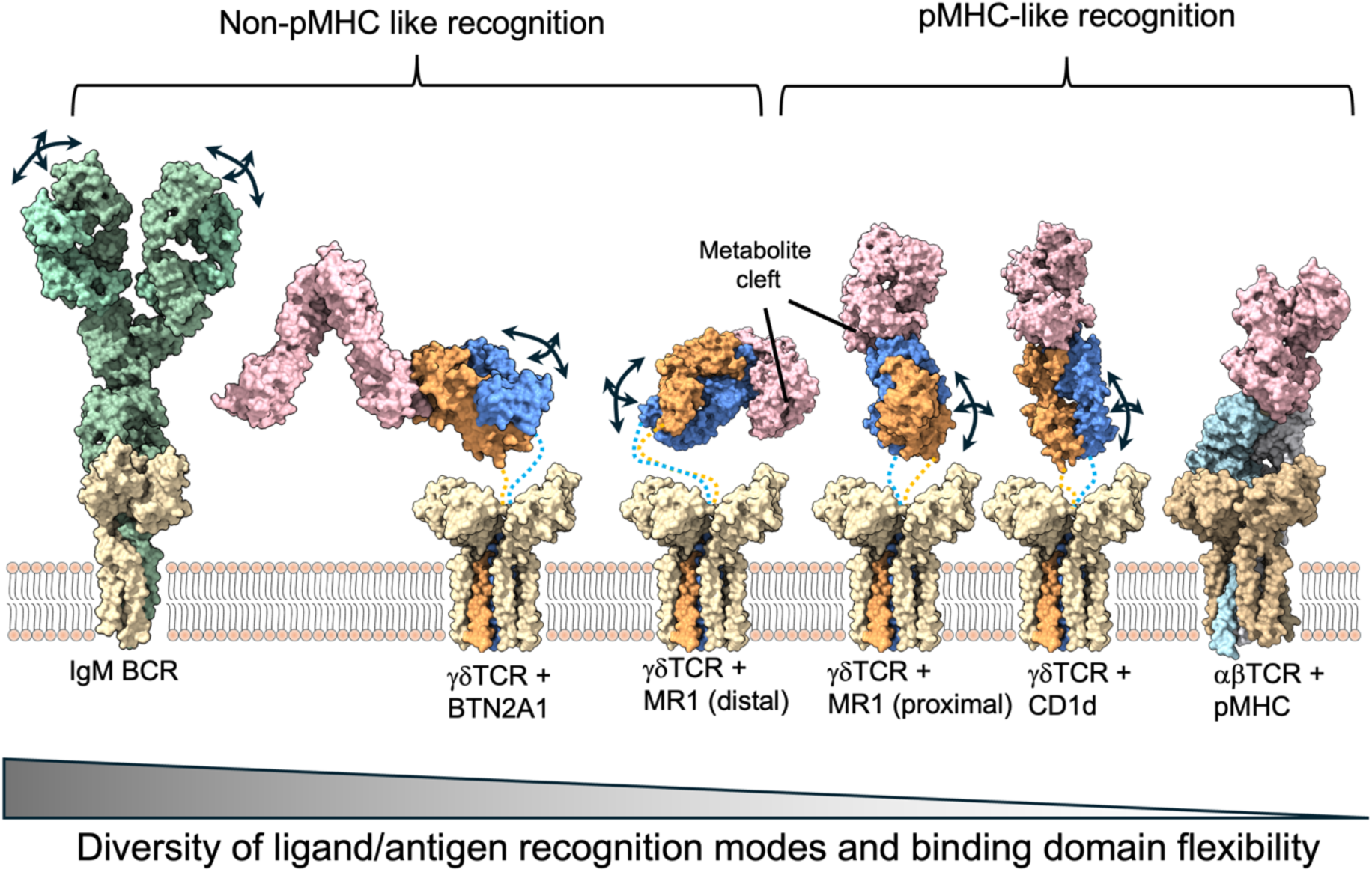
γδ TCR ECD flexibility correlates with the diversity of antigen recognition by the γδ TCR. Immune receptor models are organized by their ligand binding geometries. αβ TCR/CD3-pMHC complex (rightmost) is characterized as having a rigid ECD and conserved pMHC-docking mode. BCR (left most, IgM receptor shown) has highly mobile antigen binding (Fab) regions and unlimited ways of engaging antigens. γδTCRs (hypothetical composite models of TM/CD3 and ECD/ligand complexes shown, with flexible linkers depicted as dotted lines) have a αβTCR-like architecture but use their mobile ECDs to engage ligands in diverse ways. Arrows represent flexibility in ligand/antigen recognition domains. PDB ID 7XQ8 was used for the IgM BCR. PDB ID 8DFW was used for γδ TCR ECD/BTN2A1 complex. PDB ID 6MWR was used for the γδ TCR ECD/MR1 complex (cleft distal). PDB ID 7LLI was used for γδ TCR ECD/MR1 complex (cleft proximal). PDB ID 4LHU was used for γδ TCR ECD/CD1d complex. PDB ID 8ES8 was used for αβ TCR/CD3/pMHC complex.

**Figure S9.**
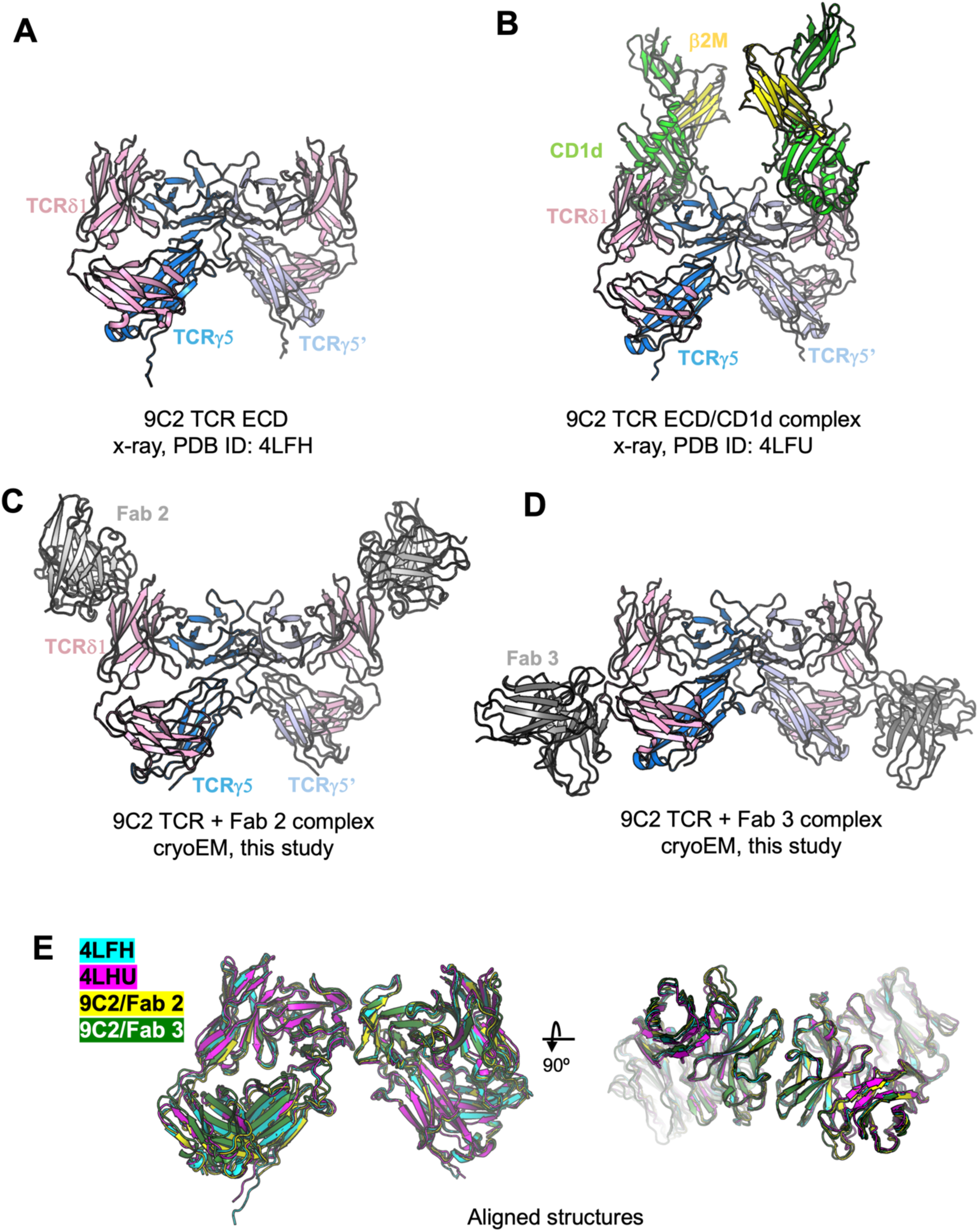
Vγ5 dimerization in 9C2 TCR ECD x-ray structures. A, B) Neighboring asymmetric units in 9C2 TCR ECD (A) and 9C2 TCR ECD in complex with CD1d (B) display a Vγ5-Vγ5 interaction. C, D) cryoEM structures of 9C2 TCR bound by Fab 2 (C) and Fab 3 (D) shown from the same viewing angle as A and B. E) Aligned structures (using TCR γ chain) from A-B with binding partners hidden for clarity. Each of the structures shows the same Vγ5-mediated dimerization mode.

